# Targeted genomic integration of EGFP under tubulin beta 3 class III promoter and mEos2 under tryptophan hydroxylase 2 promoter does not produce sufficient levels of reporter gene expression

**DOI:** 10.1101/490243

**Authors:** Aleksei G. Menzorov, Konstantin E. Orishchenko, Veniamin S. Fishman, Anastasia A. Shevtsova, Roman V. Mungalov, Inna E. Pristyazhnyuk, Elena A. Kizilova, Natalia M. Matveeva, Natalia Alenina, Michael Bader, Nikolai B. Rubtsov, Oleg L. Serov

**Affiliations:** Institute of Cytology and Genetics SB RAS, Novosibirsk, 630090, Russia; Novosibirsk State University, Novosibirsk, 630090, Russia; Immanuel Kant Baltic Federal University, Kaliningrad, 236041, Russia; Max Delbrück Center for Molecular Medicine, Berlin, 13092, Germany

**Keywords:** Mouse embryonic stem cells, tubulin beta 3 class III, Tryptophan hydroxylase 2, mEos2, neuronal differentiation, targeted genomic integration

## Abstract

Neuronal tracing is a modern technology that is based on the expression of fluorescent proteins under the control of cell type-specific promoters. However, random genomic integration of the reporter construct often leads to incorrect spatial and temporal expression of the marker protein. Targeted integration (or knock-in) of the reporter coding sequence is supposed to provide better expression control by exploiting endogenous regulatory elements. Here we describe the generation of two fluorescent reporter systems: EGFP under pan-neural marker class III β-tubulin (*Tubb3*) promoter and mEos2 under serotonergic neuron specific tryptophan hydroxylase 2 (*Tph2*) promoter. Differentiation of Tubb3-EGFP ES cells into neurons revealed that though Tubb3-positive cells express EGFP, its expression level is not sufficient for the neuronal tracing by routine fluorescent microscopy. Similarly, the expression levels of mEos2-TPH2 in differentiated ES cells was very low and could be detected only on mRNA level using PCR-based methods. Our data shows that the use of endogenous regulatory elements to control transgene expression is not always beneficial compared to random genomic integration.

## Introduction

Neuronal tracing in *in vitro* and *in vivo* models is a widely used technique in neuroscience and developmental biology. This technique allows labeling of different neuronal subtypes by fluorescent protein expression. One of the standard ways to generate animal or cell culture models for neuronal tracing is random genomic integration of an expression cassette, encoding fluorescent protein under a certain neuron-specific promoter. For instance, mice expressing YFP under tubulin beta 3 class III (*Tubb3*) promoter (Liu et al. 2007) and GFP under *Nestin* promoter (Mignone et al. 2004) are examples for such strategy. Alternatively, the DNA sequence encoding fluorescent protein can be inserted under the endogenous promoter of a gene of interest (marker gene). This approach, also called “knock-in” is becoming more popular due to the facilitation of targeted genome engineering by the use of TALENs (Cermak et al. 2011) and CRISPR/Cas systems (Jinek et al. 2012) and benefits from the accuracy of reporter expression due to the accessibility of remote regulatory elements and absence of the position effect.

There are several strategies to insert the reporter sequence under the endogenous promoter of a marker gene. First, the reporter can be placed in front of/or replacing the original coding sequence of a marker gene. An important drawback of this experimental design is that the marker protein is not expressed from the engineered allele, which may cause undesirable effects. Alternatively, both reporter and marker protein can be translated from a single chimeric RNA and be separated during translation using IRES or 2A-signal peptide. Finally, both reporter and marker protein may be produced as a fused protein. The last option ensures equimolar ratio of reporter and marker gene proteins; thereby in this system reporter expression reflects expression of the marker gene in the most precise way.

Considering advantages of targeted insertion, it may be concluded that this is a preferred type of design for reporter line generation. In this paper, we provide two examples of targeted integration of reporters under neuron-specific promoters, namely pan-neural marker *Tubb3* and serotonergic marker tryptophan hydroxylase 2 (*Tph2*). We show that reporter expression in both generated systems is weak comparing to previously published systems with random integration of the reporter.

We used two different reporters, EGFP for the panneuronal tracing; and mEos for the tracing of serotonergic cell populations. mEos2 is a photoswitchable fluorescent protein which is capable to change emission spectrum from green to red after espouse to blue (390 nm) light (McKinney et al. 2009). Eos fluorescent proteins are widely used to track intracellular distribution of peptides or organelles (Zhou and Lin, 2013). Using a construct coding for the mEos2-TPH2 fused protein we aimed to explore intracellular distribution and dynamics of the pharmacologically relevant enzyme TPH2, which is expressed in serotonergic neurons and is a rate-limiting enzyme of serotonin synthesis (Walther et al. 2003).

## Results

### Generation of Tubb3-EGFP knock-in mouse ES cell line

We produced three ES cell lines from the 129S2/SvPasCrl mouse strain: DGES1, DGES2, and DGES3. All cell lines had 40,XY karyotype, their pluripotency was assessed by teratoma formation test for DGES2 and by germ line transmission test for all cell lines (Fig. S1-S3). In all further experiments we used DGES1 as a wild type ES cell line.

To generate a Tubb3-EGFP knock-in mouse ES cell line, we designed a DNA TubbEGFPpuro construct to allow homologous recombination within the *Tubb3* gene (Fig. 1A). The 2A-EGFP and puromycin resistance cassette were flanked by homology arms to be inserted into *Tubb3* last exon replacing its stop codon, but keeping the *Tubb3* 3’-UTR intact (Fig. 1B). We introduced LoxP sites to allow removal of the puromycin resistance gene.

**Fig. 1.**
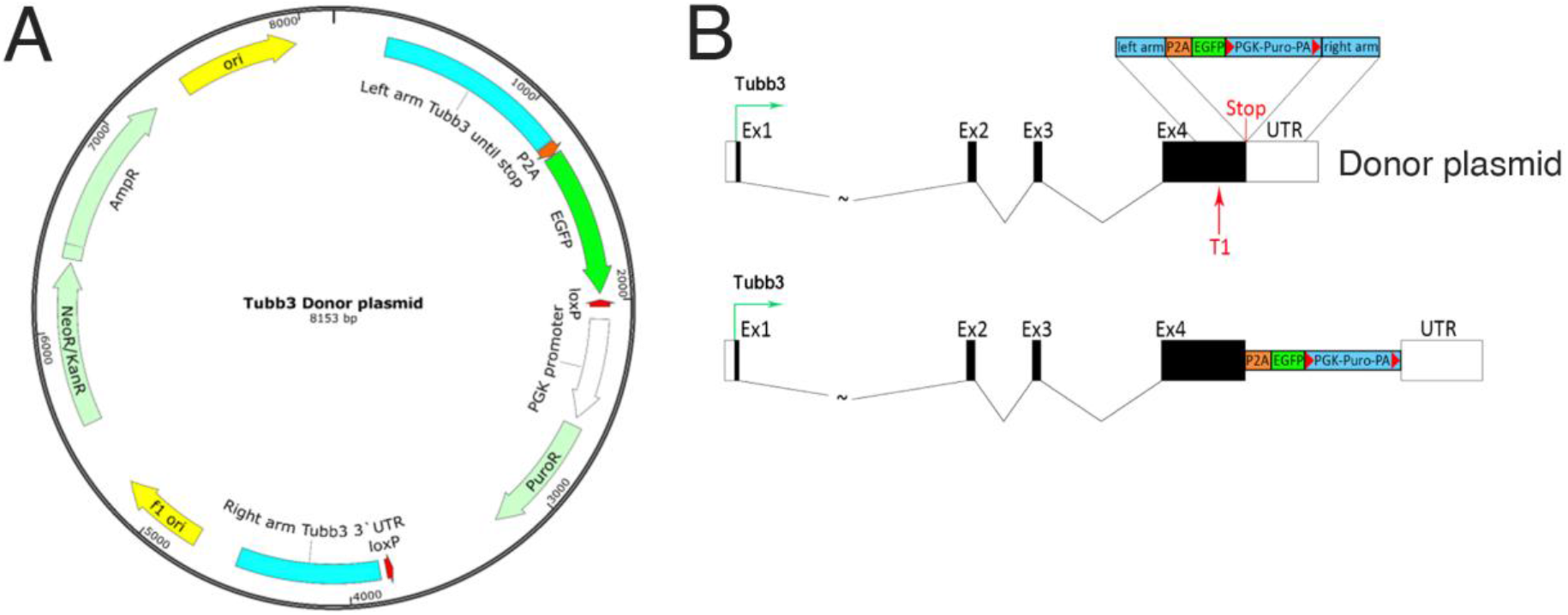
Targeting *Tubb3* locus using TALENs. **A** Map of the donor plasmid used to target the *Tubb3* locus. The 2A-EGFP and puromycin resistance cassette were flanked by homology arms to be inserted into *Tubb3* last exon replacing its stop codon, but keeping the *Tubb3* 3’-UTR intact. LoxP sites were introduced to allow removal of puromycin resistance gene; **B** Schematic overview depicting the targeting strategy for the *Tubb3* gene. The exons of the *Tubb3* locus are shown as black boxes and T1 arrow indicates the genomic site cut of the TALENs. The donor plasmid contained 5’ and 3’ homologous sequences (blue boxes) of approximately 800 bp flanking the TALEN cutting site. P2A: self-cleaving peptide sequence, EGFP: enhanced green fluorescent protein, loxP: loxP sites, PGK: human phophoglycerol kinase promoter, Puro: puromycin resistance gene, pA: polyadenylation sequence, UTR: untranslated region.

DGES1 ES cells were transfected with TubbEGFPpuro donor plasmid and TALENs to introduce DNA double strand breaks in the *Tubb3* exon 4 near the stop codon, and selected on puromycin. We produced 22 puromycin resistant clones, 2 had correct transgene integration in the genome (hemizygous) and only one of them, DGES1-TubbEGFPpuro, was diploid. To take out the puromycin-expressing cassette we transiently expressed Cre-recombinase in the DGES1-TubbEGFPpuro cells and expanded ES cell clones. The removal of an antibiotic resistance cassette was shown to positively modulate the expression of the transgene (Hockemeyer et al. 2011). Resulting clones lacking puromycin resistance were karyotyped and one DGES1-TubbEGFP clone was selected for further differentiation.

### EGFP expression in differentiated Tubb3-EGFP knock-in mouse ES cells

To evaluate if the EGFP expression is targeted to neurons, we differentiated ES cells to neuronal phenotype. As expected, EGFP colocalized with Tubb3 and with another neuronal marker, NF200, in differentiated neurons obtained from both DGES1-TubbEGFPpuro (Fig. 2A, B) and DGES1-TubbEGFP ES cells (Fig. 2C, D). Thus, we successfully produced mouse ES cell lines with Tubb3 expression marked by EGFP.

**Fig. 2.**
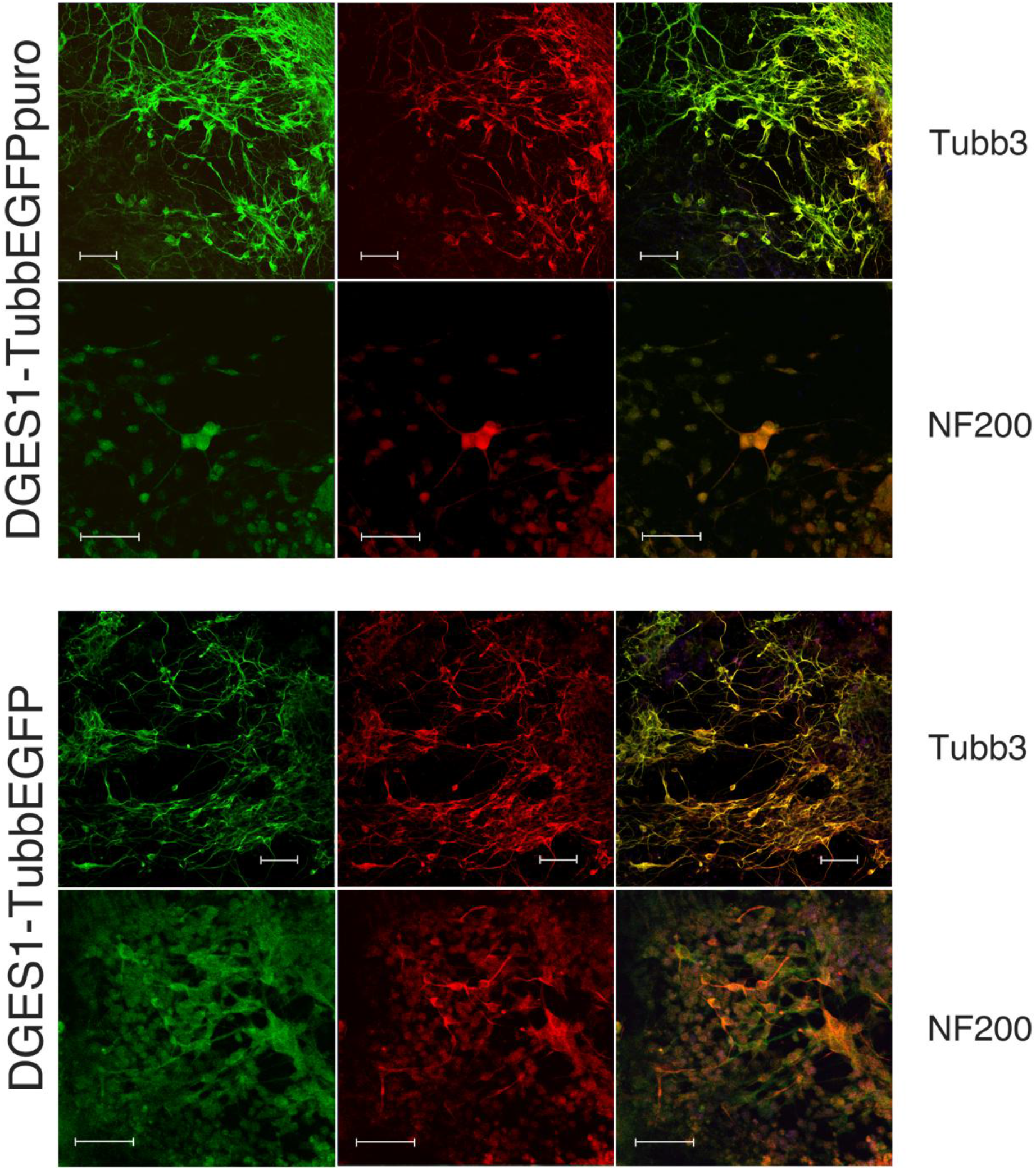
Colocalization of EGFP expression (green) and neuronal markers in Tubb3-EGFP knock-in ES cells differentiated into neurons. NF200 – immunostaining of neurofilament 200 (red), Tubb3 – immunostaining of tubulin beta 3 class III (red). Scale bars, 50 μm.

We employed flow cytometry to analyze EGFP expression in differentiated ES cells. During differentiation, only some percentage of cells became a desired cell type, neurons. Thus, we obtained two cell populations: EGFP-positive neurons and EGFP-negative undifferentiated and/or differentiated towards non-neuronal lineages cells. Flow cytometry is superior to qPCR for cell fluorescence analysis in such a case, as it measures individual cell fluorescence. An example of EGFP flow cytometry of differentiated DGES1, DGES1-TubbEGFPpuro and DGES1-TubbEGFP ES cells is presented on Figure 3A. As expected, removing the puromycin resistance gene increased EGFP expression in the resulting DGES1-TubbEGFP cell line compared to DGES1-TubbEGFPpuro (Fig. 3B). The fluorescence of DGES1-TubbEGFP is 1.63 times higher than that of DGES1-TubbEGFPpuro, *P*-value 0.013.

**Fig. 3.**
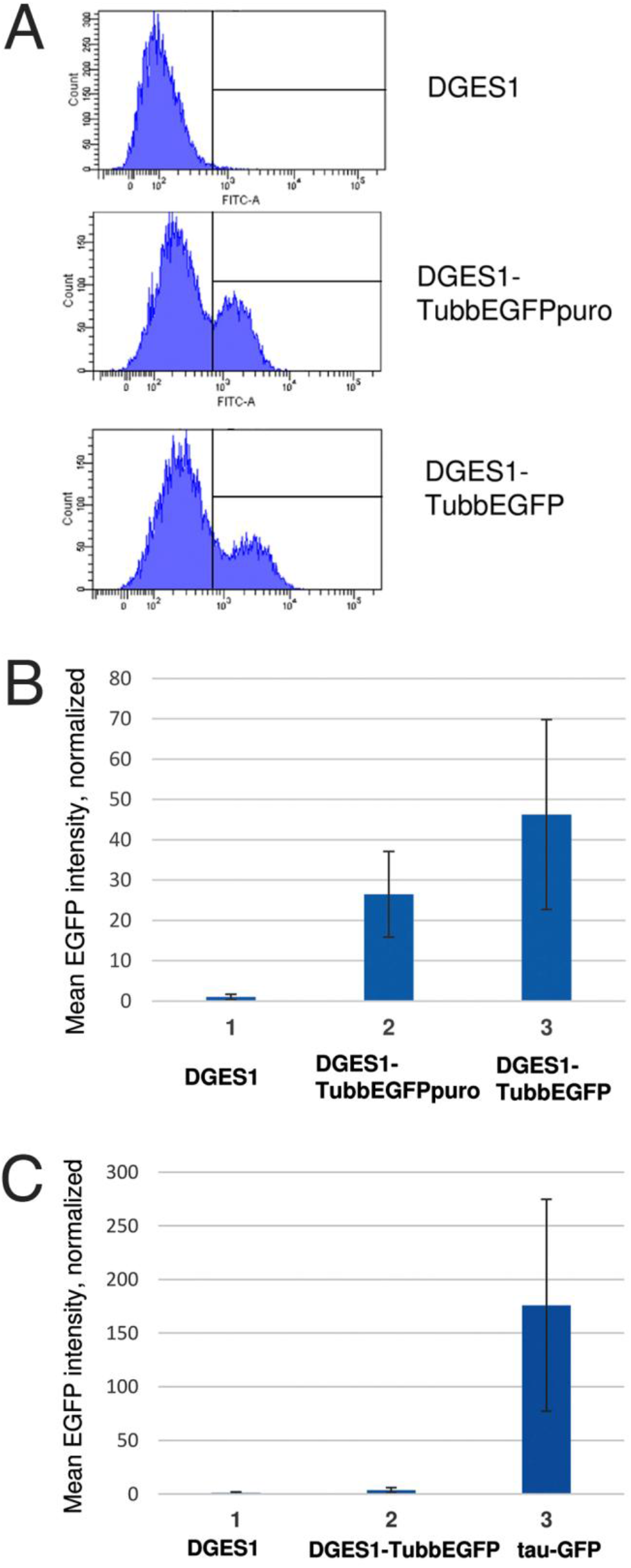
Flow cytometry analysis of EGFP fluorescence **A** EGFP fluorescence of ES cells differentiated towards neuronal phenotype; **B** Mean EGFP fluorescence in ES cells differentiated into neurons normalized to wild type cells; **C** Mean EGFP/tau-GFP fluorescence in undifferentiated ES cells normalized to wild type cells. Error bar demonstrates standard error.

We also used flow cytometry to check if EGFP expression is detectable in undifferentiated Tubb3-EGFP knock-in ES cells. It appeared that DGES1-TubbEGFP cell line has clear EGFP expression compared to DGES1 wild type cells (Fig. 3C); though the level of GFP expression was 60 times lower in comparison to E14Tg2aSc4TP6.3 ES cell line (tau-GFP), which ubiquitously expresses tau-tagged GFP (Pratt et al. 2000).

Although we were able to detect EGFP in Tubb3-EGFP knock-in ES cells differentiated to neurons by imaging and FACS, EGFP fluorescence appeared weak under the microscope. Therefore, we decided to compare differentiated ES cell fluorescence with “bright” control, Phoenix cells transfected with lentiviral vector harboring EGFP under the SFFV promoter (multiplicity of infection 2). Both flow cytometry and fluorescent microscope observation revealed significantly higher levels of EGFP in Phoenix cells, with the fluorescence intensity in Phoenix estimated to be more than 50 times higher (data not shown).

### Derivation of mEos2-TPH2 knock-in mouse ES cell line

Before producing mEos2 knock-in ES cell line we investigated whether mEos2 retains its fluorescent properties when fused to TPH2. We designed vectors expressing mEos2 fused to either N-or to C-terminus of TPH2 (hereinafter we use abbreviations mEos2-TPH2 and TPH2-mEos2 for these constructs, respectively) (Fig. 4A). For both mEos2-TPH2 and TPH2-mEos2 we observed fluorescence in the green channel when transiently transfected in PC12 cells. However, the signal was more pronounced in case of mEos2-TPH2 (Fig. 4A, B).

**Fig. 4.**
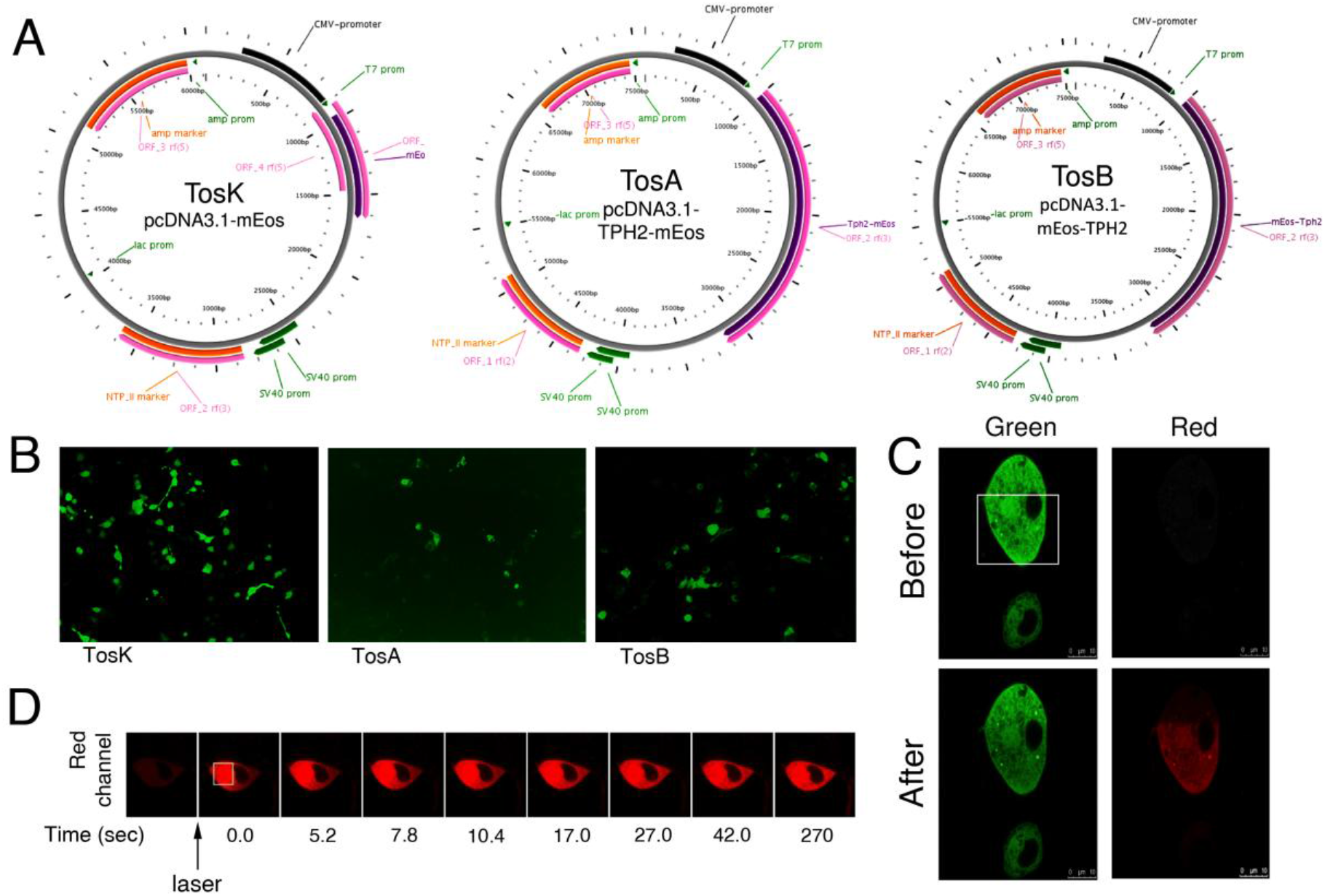
mEos2 retains its properties when fused to TPH2 protein. **A** Schematic representation of TosK, TosA, and TosB plasmids; **B** The green fluorescent signal acquired on PC12 cells transfected with TosK, TosA, or TosB plasmids; **C** Green and red fluorescent signals acquired on fixed PC12 cells before (top panel) or after (bottom panel) selected area was exposed to blue (400 nm) laser. Area exposed to laser is shown as white box; **D** Dynamic of red fluorescent signal distribution after exposure of selected area to blue (400 nm) laser in live PC12 cell. Area exposed to laser is shown as yellow box.

We were able to convert green mEos2 signal to red by applying 400 nm laser to the PC12 cells transfected with mEos2-TPH2. In formaldehyde fixed cells, the conversion occurred precisely in the area exposed to the 400 nm light (Fig. 4C). In live cells, the red signal spread over the cell cytoplasm during several seconds after conversion (Fig. 4D). Thus, mEos2 maintains its fluorescent properties when fused to the TPH2 and could be used to study intracellular localization and dynamics of the protein.

Next, we designed a vector for homologous recombination and gRNA to introduce the coding sequence of mEos2 protein upstream of the *Tph2* start codon (Fig. 5A). We co-transfected these vectors with a plasmid co-expressing Cas9 and EGFP under CMV promoter (Addgene, cat. no. 48138) into mouse DGES1 ES cells. After transfection, we sorted single EGFP-positive cells by FACS generating 94 colonies. It should be noted that EGFP expression was transient and used only to select transfected cells. The fluorescent signal was lost after several days after transfection and none of the obtained clones displayed EGFP signal. PCR analysis of obtained subclones showed that 12 of them harbored mEos2 coding sequence in their genome. Only one clone showed correct integration of the expression cassette in the *Tph2* locus. This clone was expanded to obtain DGES1-D9 cell line, which displayed typical morphology and growth rate of ES cells, as well as the expression of Oct4 and Sox2 pluripotency markers in the nuclei (Fig. 5B). We sequenced the mEos2 integration site and confirmed its correct localization.

**Fig. 5.**
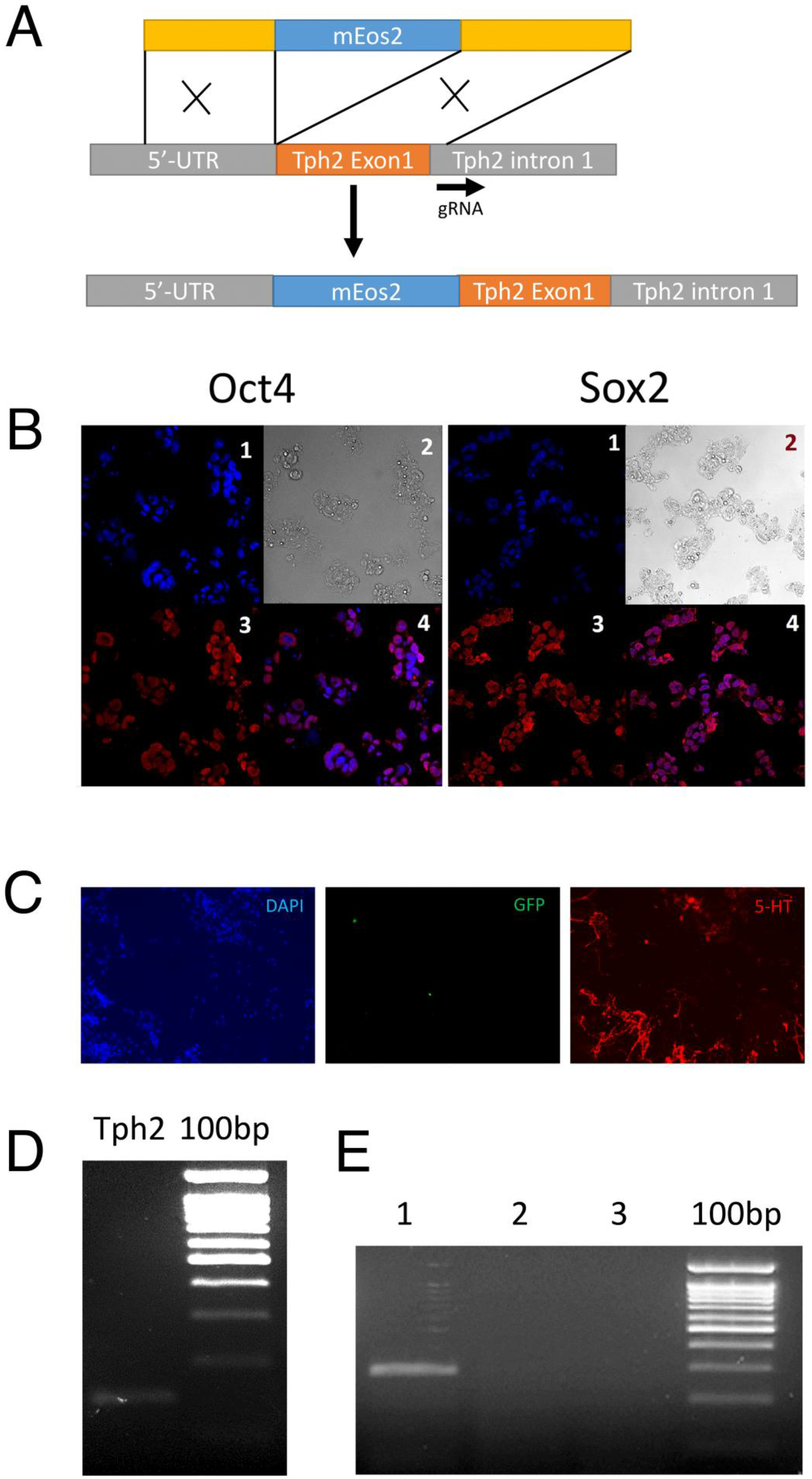
Generation and characterization of DGES1-D9 cell line. **A** Schematic representation of mEos2 knock-in strategy; **B** Immunostaining of DGES1-D9 cells for pluripotency markers Oct4 and Sox2. 1 – DAPI, 2 – Phase contrast, 3 – Oct4 (on the left panel) or Sox2 (on the right panel), 4 – signals 1 and 3 merged; **C** Immunostaining of DGES1-D9 derived neurons for serotonin; **D** Expression of *Tph2* gene in neurons obtained by differentiation of DGES1-D9 cells detected by RT-PCR (primers are specific to exons 1 and 2 of *Tph2*); **E** Expression of *mEos2-Tph2* transcript in neurons obtained by differentiation of DGES1-D9 cells detected by nested RT-PCR (primers are specific to mEos2 and *Tph2*_exon2). 1 – cDNA, 2 – RT negative control, 3 – H2O.

### Neuronal differentiation of DGES1-D9 cells and mEos2-TPH2 expression

To study expression of the mEos2-TPH2 fusion protein we induced differentiation of DGES1-D9 ES cells into serotonergic neurons using the previously published protocol (Dolmazon et al. 2011). We were able to detect a number of serotonin-positive cells with typical neuronal morphology 20-25 days after induction (Fig. 5C). However, serotonin-positive neurons were negative for mEos2 signal. We hypothesized that expression of the reporter is too weak to produce detectable amounts of mEos2-TPH2 protein. Indeed, RT-PCR analysis of differentiated cells showed expression of *Tph2* (originating from non-targeted allele), but absence of *mEos2-Tph2* transcript (Fig. 5D). We performed nested RT-PCR to trace low levels of *mEos2* expression, and found a detectable amount of a PCR product (Fig. 5E). Thus, the *mEos2-Tph2* allele was expressed in serotonergic derivatives of DGES1-D9 ES cells, but in a much less efficient manner than normal *Tph2* allele.

## Discussion

Tubb3 is a nearly perfect neuronal marker as it is expressed in neurons of the central and peripheral nervous systems. Previously, a Tubb3-reporter mouse was generated by random genomic integration of a Tubb3-YFP cassette (Liu et al. 2007). However, there were both minor spatial and temporal differences between endogenous Tubb3 expression and the YFP reporter. The reasons may be lack of necessary regulatory elements, and position effect of the transgene. Thus, it is rational to suggest that targeted integration would be beneficial for this and other reporter system. Therefore, we generated mouse ES cell line carrying EGFP inserted into the last *Tubb3* exon, replacing its stop codon but preserving the endogenous 3’-UTR. We encountered two problems: the presence of EGFP expression in undifferentiated Tubb3-EGFP knock-in cells ES cells and a relatively low EGFP expression in neurons.

EGFP expression in undifferentiated DGES1-TubbEGFP ES cells points to Tubb3 expression (Fig. 3C). There are at least two possible explanations. First, Tubb3 is indeed expressed in undifferentiated ES cells at a low level. Second, the ES cell population is not homogeneous and consists of undifferentiated Tubb3-negative cells and Tubb3-positive cells that are already committed to neuronal fate. The second explanation is corroborated by the known fact that partially differentiated cells are always present among ES cells. In our previous study we evaluated expression of *Tubb3* in undifferentiated E14Tg2aSc4TP6.3 ES cells (Matveeva et al. 2017). Actually, in our experiment, *Tubb3* was detectable in undifferentiated ES cells, and its expression level was only ten times lower than that of such pluripotency markers as *Sox2* and *Nanog*. It is still unclear, whether this *Tubb3* expression is the result of a partial differentiation or normal for undifferentiated ES cells. Most probably, undifferentiated ES cells do express *Tubb3*, as after 13 days of differentiation EGFP negative DGES1-TubbEGFP cells are overall brighter than DGES1 (data not shown). In case of a presence of differentiated cell population, we would expect the same level of fluorescence with a slight increase of brighter cells. As EGFP expression in neurons is higher than that in undifferentiated cells, background fluorescence should not be a problem.

DGES1-TubbEGFP cell line had specific EGFP fluorescence in neurons (Fig. 2). Flow cytometry allowed efficient separation of these neuronal cells. Unfortunately, EGFP signal appeared weak under the fluorescent microscope. We decided to compare EGFP expression derived from the endogenous *Tubb3-* locus in differentiated neurons with EGFP expression driven by a strong, constitutive promoter: Phoenix cells with lentiviral vector harboring EGFP under the SFFV promoter. The intensity of fluorescence in Phoenix was indeed much higher.

Thus, we produced an ES cell line that coexpressed EGFP and Tubb3 under the control of the endogenous *Tubb3* promoter; however, the level of EGFP fluorescence is not sufficient for the routine usage in fluorescence microscopy due to relatively low signal/background ratio. The previously reported random genomic integration of an *YFP* under *Tubb3* promoter produced better results, though the transgene expression pattern did not precisely match Tubb3 expression (Liu et al. 2007). Nevertheless our cells are suitable to select neurons by flow cytometry.

As a second approach we have created a new reporter mouse ES cell line DGES1-D9 that harbors mEos2-TPH2 fused protein under endogenous *Tph2* promoter. These cells were differentiated into serotonergic neurons and had correct *mEos2* transgene expression at a very low level, detectable by nested RT-PCR. This cell line turned out not to be suitable to study TPH2 trafficking, as mEos2 was not detectable by fluorescence microscopy or flow cytometry.

There are several possible approaches to increase the level of the fluorescent signal. For instance, a brighter fluorescent protein could be used. Another option is to utilize a signal amplification system such as SunTag (Tanenbaum et al. 2014). A repeating peptide array is added to the gene of interest, and up to 24 copies of antibody-fusion GFP could be recruited to it. The technical problem of that system is that it requires two separate genomic integrations. The transgene position, before or after the gene’s coding sequence, may also influence the level of expression.

## Conclusions

Random genomic integration of a fluorescent protein under the cell type-specific promoter is widely used in gene expression studies. The main drawback of that approach is that the regulation of the fluorescent protein expression may differ from that of a gene of interest as some regulatory elements are absent, and due to the position effects of the transgene or its silencing as a result of multiple-copy integration. Insertion of a fluorescent protein coding sequence into the beginning or the end of the coding frame of a specific gene allows to retain the gene expression pattern by exploiting endogenous regulation. We produced two knock-in mouse ES cell lines that expressed fluorescent proteins under the control of *Tubb3* and *Tph2* endogenous promoters. In both cases the expression of a fluorescent protein was not strong enough for routine usage in fluorescent microscopy. Our data shows that knock-in reporter systems are not always preferable to random genomic integration.

## Methods

All animal studies were undertaken with prior approval from Interinstitutional Bioethical Committee of ICG SB RAS. The mice were kept in a 12-hour light/dark cycle, with controlled humidity and temperature environment and fed *ad libitum.*

### Derivation and characterization of mouse ES cells

We produced mouse ES cell line DGES1 from 129S2/SvPasCrl mouse strain using the protocol developed by Brija et al. (2006). Mice were provided by the Center for Genetic Resources of Laboratory Animals (RFMEFI61914X0005, RFMEFI62114X0010) at ICG SB RAS. Cell culture was performed as described previously (Menzorov et al. 2016) with a minor modification: cells were cultured in ES cell medium containing 7.5% ES cell qualified FBS (Thermo Fisher Scientific, USA) and 7.5% KSR (Thermo Fisher Scientific, USA). ES cell lines are available at the Collective Center of ICG SB RAS “Collection of Pluripotent Human and Mammalian Cell Cultures for Biological and Biomedical Research” (http://ckp.icgen.ru/cells/; http://www.biores.cytogen.ru/icg_sb_ras_cell/).

### ES cell cytogenetic analysis

Cytogenetic analysis for ES cell lines was carried out during passages 6– 10. Preparation of metaphase chromosomes from the cells was performed as previously described (Matveeva, Fishman, Zakharova, Shevchenko, Pristyazhnyuk, Menzorov and Serov, 2017). Metaphase plates were analyzed using a Carl Zeiss Axioplan 2 imaging microscope (Jena, Germany) with CoolCube1 CCD-camera and processed using the ISIS (In Situ Imaging System, MetaSystems GmbH, Altlussheim, Germany) software in the Public Center for Microscopy SB RAS, Novosibirsk. 40 metaphase plates were counted for a cell line. Karyotyping was performed on 10 metaphase plates.

### Generation and analysis of teratomas

For teratoma formation, we used SCID hairless outbred mice (Crl:SHO-PrkdcscidHrhr) of SPF status. Experiments were performed in the Center for Genetic Resources of Laboratory Animals (RFMEFI61914X0005, RFMEFI62114X0010) at ICG SB RAS. The protocol was described earlier (Menzorov, Pristyazhnyuk, Kizilova, Yunusova, Battulin, Zhelezova, Golubitsa and Serov, 2016).

### Generation and analysis of chimeric mice

The protocol was described earlier (Menzorov, Pristyazhnyuk, Kizilova, Yunusova, Battulin, Zhelezova, Golubitsa and Serov, 2016). The experiments were performed at the Common Use Center Vivarium for Conventional Animals.

### ES cell differentiation

DGES1, DGES1-TubbEGFPpuro and DGES1-TubbEGFP ES cell lines were differentiated into neurons using a protocol (Fico et al. 2008) with a number of cells per cm^2^ 50.000-75.000. Differentiation of DGES1-D9 ES cells was performed according to previously published protocol (Dolmazon, Alenina, Markossian, Mancip, van de Vrede, Fontaine, Dehay, Kennedy, Bader, Savatier and Bernat, 2011).

### DNA transfection

We used Lipofectamine 3000 Reagent (Thermo Fisher Scientific, USA) to transfect mouse ES cells according to manufacturer’s recommendations.

### Lentiviral transduction

To introduce EGFP into Phoenix cell line we used LeGO-G2 vector (Addgene, 25917) according to previously described protocol (Menzorov et al. 2015).

### Immunofluorescence analysis

The immunofluorescence analysis was performed as previously described (Fishman et al. 2015). We used the following antibodies: rabbit anti-neurofilament 200 (Sigma, cat. no. N4142), mouse anti-neuronal class III b-tubulin (Covance, cat. no. MMS-435P), rabbit anti-serotonin (Sigma, cat. no. S5545), AlexaFluor 546 rabbit anti mouse IgG (H+L) (Life Technologies, cat. no. A11060), AlexaFluor 546 goat anti rabbit IgG (H+L) (Life technologies, cat. no. A11010), AlexaFluor 488 goat anti rabbit IgG (H+L) (Life Technologies, cat. no. A-11008), AlexaFluor 488 goat anti mouse IgG (H+L) (Life Technologies, cat. no. A-11001). Images were examined on a LSM 510 META Laser Scanning Microscope (Zeiss) on the base of AxioImager Z1 (Zeiss) microscope (Jena, Germany) with AxioCam MRm CCD-camera or on a SP5 microscope (Leica, Germany) and processed, using the AxioVision LSM510 software in the Public Center for Microscopy SB RAS, Novosibirsk or ImageJ version 1.48v (Schneider et al. 2012).

### Flow cytometry analysis

Analysis of cells was conducted on a FACSAria flow cytometer (BD Bioscience) at the Flow Cytometry Center for Collective Use at the Institute of Cytology and Genetics SB RAS. The comparison between cell fluorescence intensity (I) was calculated as follows: I_final_ = (I_experiment_ – I_control_) / I_control_.

### RNA isolation, cDNA synthesis, and RT-PCR

RNA was extracted using TRIzol reagent (Thermo Fisher Scientific) according to manufacture instructions. After DNase I treatment, RNA was subjected to reverse transcription and first-strand synthesis using RevertAid Reverse (Thermo Fisher Scientific) with random hexamers according to manufacture instructions. Primers for RT-PCR were the following: mEos2_F 5’-ACAACAAGGTTAAGCTGTATGAGC-3’, Tph2Exon2_Router 5’-ACCAG-CCCACCAACTTCATT-3’ and Tph2Exon2_Rinner 5’-AGGAGAACACTACCGCTGTC-3’ for mEos2-Tph2 nested PCR; seqTph2Exon1_F 5’-TTCTGCTGTGCCAGAAGATCA-3’ and Tph2Exon2_Rinner for Tph2 PCR.

### Design of DNA constructions, TALENs and gRNA

The donor plasmid was designed to integrate a P2A-eGFP-Puro cassette instead of the Tubb3 gene stop codon. The vector OCT4-2A-eGFP-PGK-Puro was a gift from Rudolf Jaenisch (Hockemeyer, Wang, Kiani, Lai, Gao, Cassady, Cost, Zhang, Santiago, Miller, Zeitler, Cherone, Meng, Hinkley, Rebar, Gregory, Urnov and Jaenisch, 2011). For the generation of the donor plasmid sequences up- and downstream of the stop codon were amplified using genomic DNA with the following primers: for left homology arm (949 bp) – Tubb3-Bam 5’-TCACGGATCCGGGGCACAGGCTCAGGCATG-3’ and Tubb3-Nhe 5’-AGCAGCTAGCCTTGGGCCCCTGGGCTTCCG-3’, for right homology arm (721 bp) – Tubb3-Asc 5’-GGGCGGCGCGCCAGTTGCTCGCAGCTGGGGTG-3’ and Tubb3-Not 5’-GTAGGCGGCCGCGGAAGAATGCTGGATATGAG-3’. Amplified arms were cloned in the OCT4-2A-eGFP-PGK-Puro vector by BamHI/NheI and AscI/NotI sites for left and right homology arms respectively.

TALENs of 20 RVDs each were designed near the stop codon of the Tubb3 gene to bind to the following sequences: 1L 5’-GTCCGAGTACCAGCAGTACC-3’ and 1R 5’-CATACATCTCCCCCTCCTCC-3’. TALEN constructions were created using the Golden Gate Assembly. The plasmid kit used for generation of TALENs was a gift from Daniel Voytas and Adam Bogdanove (Addgene kit #1000000024) (Cermak, Doyle, Christian, Wang, Zhang, Schmidt, Baller, Somia, Bogdanove and Voytas, 2011).

Cre-expressing plasmid was created by assembling CMV-promoter, Cre-2A-GFP coding sequence, SV40-polyA signal and pGEM-T backbone (Promega).

Plasmids for transfection were extracted with PureYield™ Plasmid Maxiprep System (Promega) according to the manufacturer’s recommendations.

Screening ES cell clones for a correct transgene integration site in *Tubb3* was performed by PCR with the following primers: Tubb3_LS-f 5’-AGATGTCGTGCGGAAAGAGT-3’ and Tubb3_LS-r 5’-GAACTTCAGGGTCAGCTTGC-3’ for the left border and Tubb3_RS-f 5’-GCCTGAAGAACGAGATCAGC-3’ and Tubb3_RS-r 5’-ATGGAGCCAGTACAGGGTTG-3’ for the right border of the transgene integration. To determine whether transgene is hemi-or homozygous the following primers were used: Tubb3-5F 5’-GTCAAGGTAGCCGTGTGTGA-3’ and Tubb3-3R 5’-TCTCCAATACCAGGCAGAGG-3’.

To generate mEos2 knock-in we used a donor plasmid containing mEos2 sequence flanked by 5’- and 3’-homology arms (~500 bp) and the following gRNA: 5’-CTCTACAGCAGGTGTCCATA-3’ (PAM sequence TGG). Homology arms of donor plasmid correspond to 5’-UTR and first intron of *Tph2* gene. For genotyping, we used primers located outside of homology arms with primers located within *mEos2* sequence: Tph2_5UTR_F 5’-CCATGGGTTTTGGAAGCAGAC-3’ and mEos2_R 5’-AGGCTTGCCTGTACCATCTC-3’; Tph2_intron1_R 5’-TCTTGTAAATATTGAAGCAAACAGA-3’ and mEos2_F.

## Supporting information

## Authors’ contributions

AGM, KEO, VSF, AAS, RVM, EAK, NMM, NA, MB, NBR, OLS

AGM produced mouse ES cell lines, produced Tubb3-EGFP knock-in cell lines, performed their differentiation, provided analysis and interpretations of the data and together with KEO and VSF is the principal investigator of the project; KEO produced Tubb3-EGFP vectors and performed PCR analysis of the cell lines; VSF produced mEos2-TPH2 vectors, produced mEos2-TPH2 knock-in cell line, performed its differentiation and analyses; AAS produced mouse ES cell lines, generated Tubb3-EGFP knock-in cell lines and performed their differentiation; RVM performed RT-PCR of mEos2-TPH2 knock-in cell line; IEP carried out cytogenetic and immunocytochemical analyses; EAK carried out work with animals and teratoma histochemical analysis; NMM produced mouse ES cell lines; NA and MB carried out interpretations of the data and mEos2 project coordination; NBR and OLS carried out interpretations of the data and Tubb3 project coordination; AGM and VSF did most of the writing with contributions from all authors. All authors read and approved the final manuscript.

## Acknowledgements

The study was supported by the state project No. 0324-2018-0016 of the Ministry of Education and Science of Russia. Cell lines are available at the Collective Center of ICG SB RAS “Collection of Pluripotent Human and Mammalian Cell Cultures for Biological and Biomedical Research” (http://ckp.icgen.ru/cells/; http://www.biores.cytogen.ru/icg_sb_ras_cell/).

