## Supplementary material for "Targeted genomic integration of EGFP under tubulin beta 3 class III promoter and mEos2 under tryptophan hydroxylase 2 promoter does not produce sufficient levels of reporter gene expression"

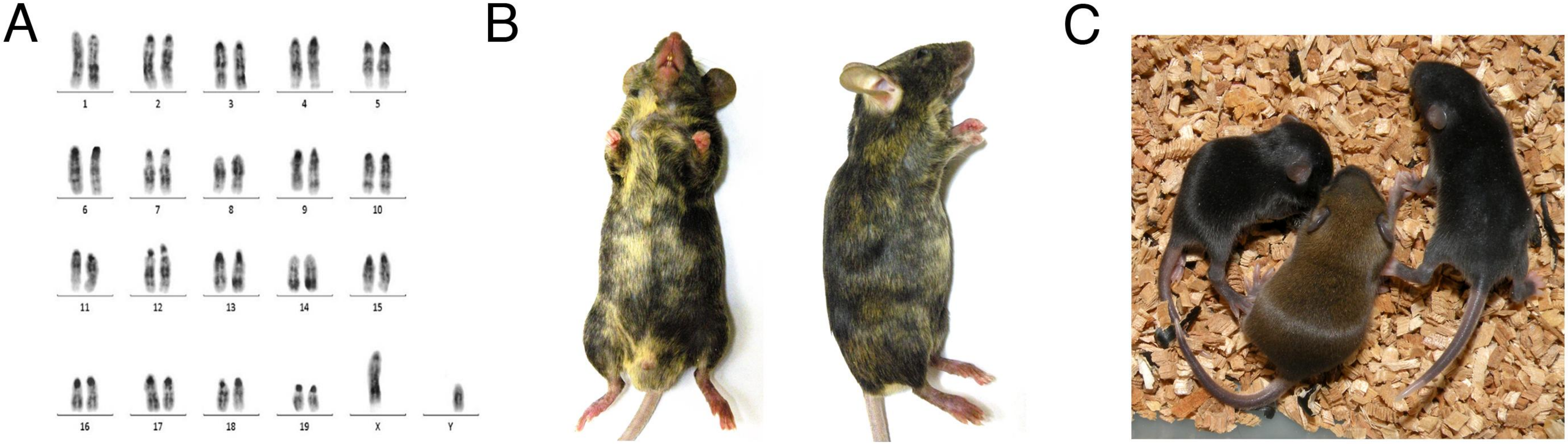


**Fig. S1** Characterization of DGES1 ES cell line. **A** Karyotype; **B** Chimeras on C57BL/6J background; **C** Germ line transmission (breeding to C57BL/6J background).

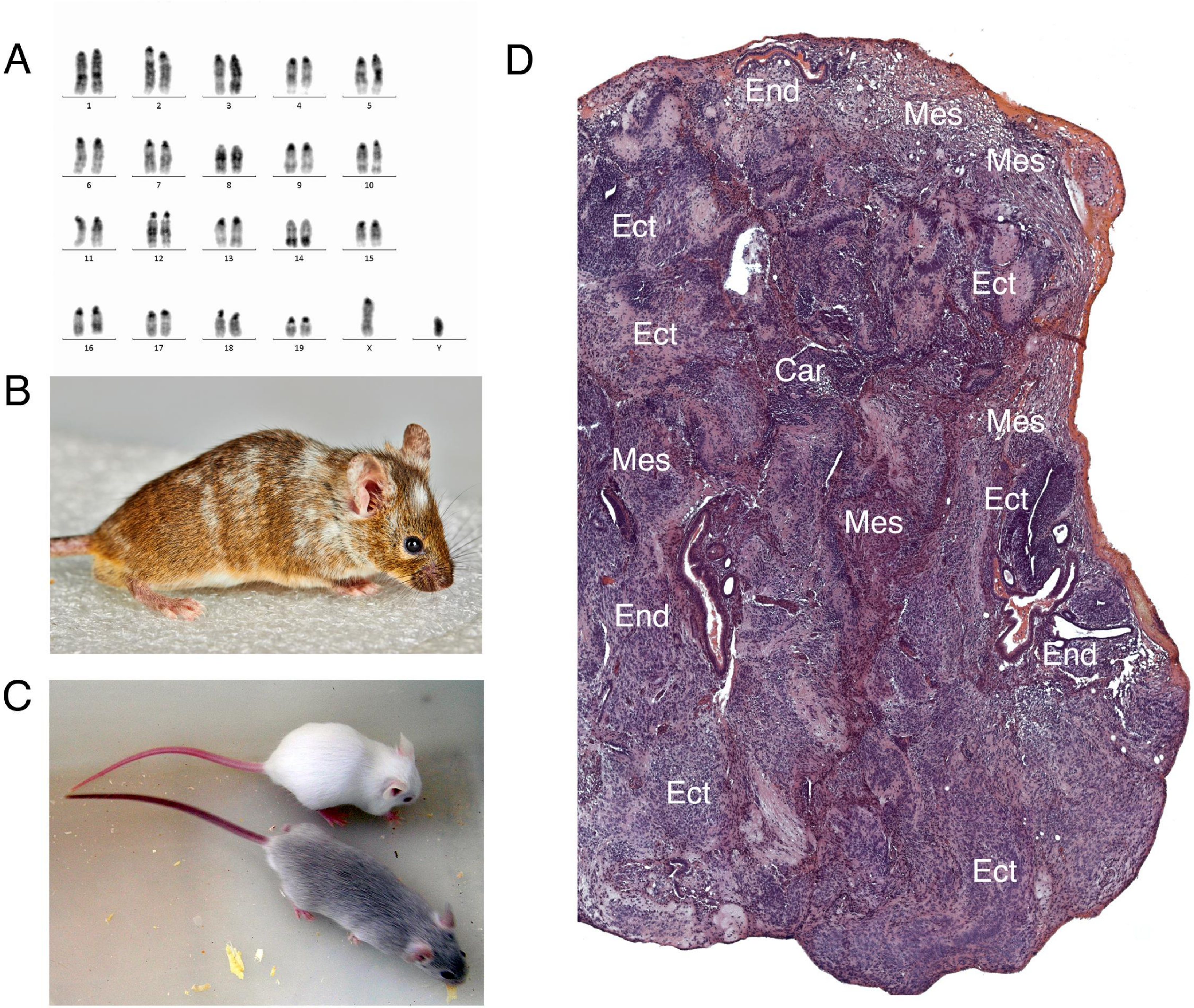


**Fig. S2** Characterization of DGES2 ES cell line. **A** Karyotype; **B** Chimera on CD-1 background; **C** Germ line transmission (breeding to CD-1 background); **D** Teratoma formation in SCID (SHO-PrkdcscidHrhr) mice.

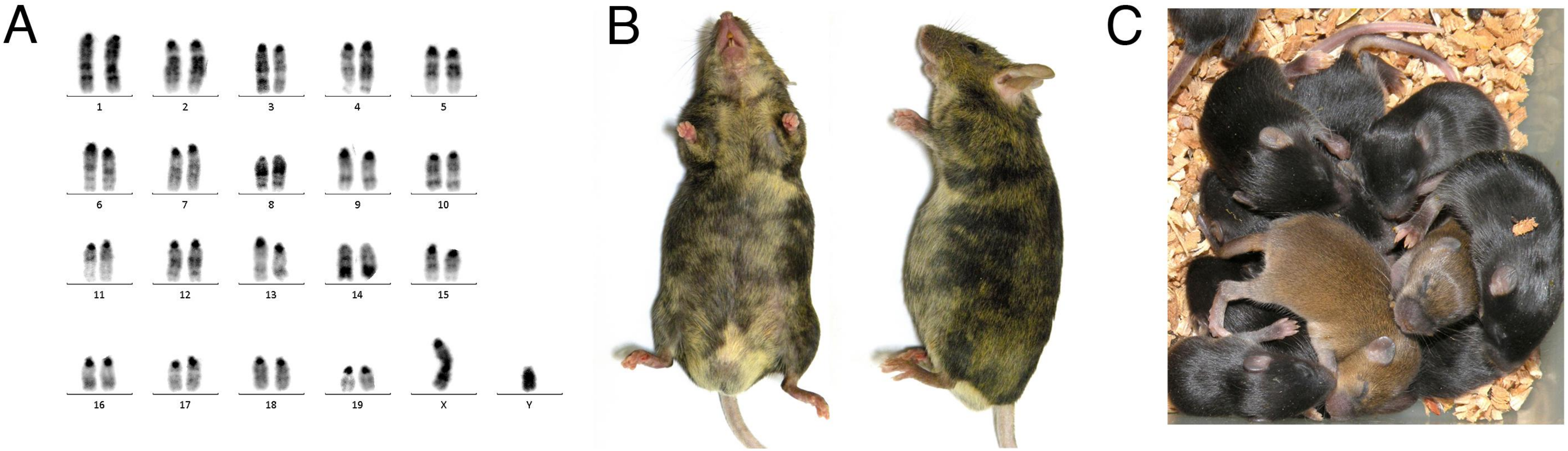


**Fig. S3** Characterization of DGES3 ES cell line. **A** Karyotype; **B** Chimeras on C57BL/6J background; **C** Germ line transmission (breeding to C57BL/6J background).
